# Population-Scale Landscape of Mobile Element Insertions Across 82 Diverse Indian Populations

**DOI:** 10.64898/2026.07.29.741658

**Authors:** Sagar S, Shreya Ganjigatte Srinivas, Ruchitha Gangaiah Byrohalli, Vidushi Dwivedi, Hiba Ali, Khader Valli Rupangudi, Ramesh Yelagandula, GenomeIndia Consortium, Divya Tej Sowpati, Ishaan Gupta, Pankaj Yadav, Shweta Ramdas

## Abstract

Mobile element insertions (MEIs) are a major source of structural variation in human DNA that can affect gene regulation and disrupt coding sequences. Population-scale MEI catalogs remain disproportionately skewed toward European-ancestry cohorts, with limited representation from India despite comprising nearly one-sixth of the global population and encompassing thousands of genetically distinct, largely endogamous communities. We characterized non-reference Alu and LINE1 insertions in 7,478 whole genomes from the GenomeIndia project, representing 82 populations selected to capture India’s geographic, linguistic, and social diversity. We cataloged 26,977 Alu and 7,075 LINE1 insertions, identifying 18,982 previously unreported insertions absent from existing publicly available MEI databases. Insertions bore signatures of purifying selection, depleted in exons and active chromatin and enriched in quiescent regions. MEI variants recapitulated the population structure seen in SNP data. Regulatory variation associated with MEIs was strongly population-structured, with a substantial fraction of European-defined MEI-eQTLs absent in the GenomeIndia cohort or carried at sharply different frequencies across communities, and MEIs in high linkage disequilibrium with GWAS variants. Sixty-nine insertions fell within exons, including a truncating LINE1 in *CYP2J2* carried across five populations and a community-specific Alu in *CABYR* with a founder-effect signature. Collectively, these results reveal a layer of population-specific mobile element variation in Indian populations that is not captured by existing European-anchored reference resources. These variants represent potentially novel sources of genetic variation contributing to lack of transferability of disease associations across populations.

## Introduction

Mobile genetic elements make up more than half of the human genome, yet their contribution to genome function remains comparatively understudied. Once viewed largely as selfish DNA in an evolutionary arms race with the host, transposable elements (TEs) are now recognized as substrates for genome function^1^. They are co-opted into regulatory roles across development and disease^2^, with human-specific TEs enriched in enhancers active in the brain and hippocampus^3,4^. Because TE activity and the regulatory elements they seed can differ across human populations, the functional consequences of TE variation are also expected to be population-specific. Yet these effects remain largely uncharted.

This gap is partly driven by technical limitations. Mobile element insertions (MEIs) are not captured by standard variant-calling pipelines such as GATK, and their length and repetitive sequence composition make them error-prone to call from short read sequencing data^5,6^. Accurate MEI discovery therefore demands specialized tools and stringent quality control. As a result, population-scale MEI studies remain scarce even as SNP- and indel-based studies have proliferated^7^. This shortfall is most acute in under-represented populations: the well-documented Eurocentric bias of human genetics is magnified for structural variants, where reference resources are the most sparse.

India, with roughly one-sixth of the world’s population, and structured into more than 4,000 largely endogamous communities with distinct genetic pools, is among the most consequential of these gaps. The largest prior MEI characterizations of Indian individuals, from the 1000 Genomes and IndiGen projects, each sampled about 1,000 individuals (IndiGen across 25 major ethnic groups) representing only a narrow slice of this genetic diversity^8–10^. Here, we characterize MEIs in 7,478 whole genomes from the GenomeIndia (GI) project, representing 82 populations selected to span the major axes of geographic, linguistic, and social variation across India^11,12^. Benchmarking against long-read sequencing, we catalog Alu and LINE1 insertions, map their distribution across functional and regulatory genome annotations, and identify exonic and regulatory MEIs that are enriched in, or private to, specific Indian populations. Together, these analyses constitute the largest and most diverse survey of mobile-element variation in an Indian cohort to date, providing a view of how this variation is structured across India’s diverse populations.

## Methods

### Sample information

The GI dataset comprises whole-genome sequences from 9,768 individuals representing 83 diverse ethnic populations, including geographically isolated and previously unstudied tribal groups. A granular breakdown of the sample collection, sequencing workflow, quality metrics is detailed in the GI flagship publication^11,12^. This study focuses on a core subset of 7,478 individuals available at the time of analysis, representing 82 distinct populations (Table S1). Long-read sequencing data from 16 of these samples were additionally used for validation.

### Variant calling

We called mobile element variants using the Mobile Element Locator Tool (MELT) v2.2.2^13^. We identified non-reference Alu and LINE1 insertions using the MELT-SPLIT runtime without utilizing a priors’ list, allowing de novo discovery of insertion events. We filtered raw MELT calls using BCFtools v1.21^14^ to remove calls failing MELT’s “PASS” filter or with an ASSESS score less than 5. Genotypes annotated as both an Alu and a LINE1 insertion in the same sample were considered ambiguous and excluded by assigning a missing genotype (./.) for those samples at the affected loci.

Although MELT-Deletion calls are not included in the Results, a short note detailing the landscape of deletions in GI can be found in Supplemental Text S1.

### Validation using long-read sequencing

We used the Oxford Nanopore PRO114_DNA_e8_2_400K:FLO-PRO114M:SQK-LSK114:400 protocol to obtain long-read sequencing data for a subset of 16 samples from the GI dataset: nine selected at random, and seven selected as suspected carriers of an exonic insertion (see results). We performed basecalling using Dorado v0.5.2^15^, followed by alignment to the hg38 reference. Samples had a mean coverage of 30X.

We used xTea_long_release_v0.1.0^16^ and Sniffles v2.3.3^17^ to identify non-reference MEIs and reference TE deletions in ONT long reads and to obtain the genotype information. We conservatively retained insertions independently reported by both xTea and Sniffles, merging variants within 20 bases of each other. This consensus call set was subsequently used to benchmark MEI calls generated by MELT.

### Comparison with existing MEI databases

To identify which of our MEIs were unique to GI, we compared our callset to eight other MEI databases (Table 1). To account for positional differences in insertions called by different methods and different versions of MELT, we tested overlap using progressively increasing positional tolerance windows between GI and other databases. We allowed a ±20 bp positional tolerance for insertion sites as overlap did not substantially increase beyond that window (Figure S1). All variants labelled as Alu or LINE1 MEIs in other databases were retained for comparison as insertion variants.

**Table 1.** Comparison of GI MEIs with other MEI databases. Mean refers to the mean number of non-reference variants carried by an individual

| Database | Samples (Coverage) | Number of Variants (Mean<sup>a</sup>) |  | Overlap with GI |  |
| --- | --- | --- | --- | --- | --- |
|  |  | Alu | LINE1 | Alu | LINE1 |
| GI | 7,478 (~37X) | 26,977 (944) | 7,075 (108) | - | - |
| 1KGP Phase 3 SVs<sup>9</sup> | 2,504 (7.4X) | 12,743 (897) | 3,047 (128) | 5,449 | 917 |
| 1KGP Phase 4 SVs<sup>10</sup> | 3202 (30X) | 29,036 (903) | 2,967 (79) | 7,447 | 797 |
| 1KGP ONT-Vienna long reads SVs<sup>42</sup> | 967 (16.7X) | 23,084 (1335) | 4,446 (187) | 6,244 | 1,001 |
| gnomAD-SV (v4)<sup>43</sup> | 63,046 (32X) | 173,374 (-) | 30,223 (-) | 8,670 | 1,553 |
| SGDP<sup>44</sup> | 296 (43X) | 11,661 (832) | 1,886 (98) | 5,944 | 880 |
| nstd211<sup>45</sup> | 3,202 (30X) | 41,254 (1685: 2,504 samples) (1,531: 698 samples) | 10,956 (297: 2,504 samples) (193: 698 samples) | 8,934 | 1,728 |
| HMEID<sup>46</sup> | 2,998 (26.2X NyuWa)<br>2,677 (7.4X 1KGP) | 26,553 (1,035:NyuWa) (884: 1KGP) | 7,353 (145:NyuWa) (119: 1KGP) | 7,677 | 1,300 |
| IndiGen (full callset)<sup>8</sup> | 1,021 (25-30X) | 21,981 (1680.62) | N/A | 7,050 | N/A |

a. Per-individual means reflect each study's caller, version, and filtering thresholds and are not directly comparable across rows. The IndiGen row is the full callset prior to PASS/ASSESS/HWE filtering, used here so that the novelty comparison is conservative; the published quality-filtered IndiGen map comprises 9,239 Alu sites at a mean of 770 per individual.

### Functional Annotation

We annotated insertions with the genomic regions within which they were present using ANNOVAR (v2025-03-02)^18^, using Ensembl gene annotations (GENCODE V46). For each MEI, we extracted the position information from MELT’s output VCF and converted them to an avinput format, as required by ANNOVAR. We allowed for a splice threshold of 5 bp, to identify MEIs close to a splice site.

To identify the regulatory regions within which variants were present, we annotated them using the chromatin state annotations provided by Epigenomics Roadmap^19^ across 127 reference epigenomes (15-state core model, hg38 dense .bed format). An MEI was assigned to a regulatory state if its insertion coordinate overlapped a Roadmap chromatin state annotation by at least 1 bp.

To check for preferential enrichment or depletion of MEIs within specific genomic regions and epigenomic states, we performed enrichment analysis using Fisher’s exact test for each of the 15 core ChromHMM chromatin states and 5 broader functional categories (active regions, repressed regions, transcribed regions, enhancers, and promoters), as well as 12 ANNOVAR categories. We computed odds ratios with 95% confidence intervals and p-values to quantify the direction and significance of enrichment.

For each tissue and chromatin state (or category), we merged the annotated genomic intervals for that state and calculated total base-pair coverage within and outside it. We intersected Alu and LINE1 insertion sites with these annotations using ‘bedtools intersect’ to determine the number of insertions falling within and outside a state. A 2×2 contingency table was used to compare the number of insertions located within and outside the state against the corresponding genomic bp coverage inside and outside that region. This approach was used to test whether MEIs occurred more (enriched) or less (depleted) than expected within a specific state or genomic annotation relative to its genomic size.

We also performed enrichment analysis using the same method to determine whether MEIs were enriched or depleted in early- and late-replicating domains (ERD and LRD). The annotations for ERD and LRD were obtained from the replication timing analysis reported by Liu *et al.*, 2016^20^. Specifically, the genomic segmentation data were taken from the supplementary file *GSE53984_GSM923453_Bg02es_Rep1_segments.bed.gz*, available on the Gene Expression Omnibus under accession GSE53984. For this analysis, population-group-specific Alu insertions, defined here as non-singleton Alu MEIs observed exclusively within each of the five major GI population groups (see results), were separately tested for enrichment.

### Linkage disequilibrium with GWAS SNPs and eQTLs

To understand how our MEIs acted as modifiers of disease risk, we first estimated linkage disequilibrium (LD) between common MEIs in our dataset (MAF≥0.01) and common SNPs from the same individuals from the pooled GI dataset. After retaining MEI-SNP pairs that had a high LD (r^2^>0.7), we narrowed down to MEIs in high LD with SNPs which were also present in the NHGRI-EBI catalog of human genome-wide association studies (all associations v1.0, accessed on 5th March, 2026)^21^, and those present as significant eQTLS (‘qval’ ≤ 0.05) in the adult GTEx single-tissue cis-eQTL data (V11, accessed on 22nd June, 2026) ^22^. We used PLINK version 1.9^23^ to estimate LD and PLINK version 2^23^ for SNP pruning and variant filtering.

We used previously published eQTL resources for MEIs from GTEx ^24^ and 1000Genomes LCLs^25^. Overlaps between these datasets and our MEI calls were made by overlapping genomic coordinates of the insertion with eQTLs within 20bp upstream or downstream of the MEI in hg38.

### Founder haplotype analysis of the CABYR insertion

To characterize the haplotype background of the *CABYR* (OMIM:612135)^26^ AluYa4 insertion (chr18:24,159,972) and test for evidence of a shared founder haplotype among carriers, we jointly phased the merged chr18 GI SNP + MEI VCF with ShapeIt4.2^27^, using the corresponding GRCh38 genetic map. From the phased VCF we used BCFtools-1.21 to extract the ±500 kb window centered on the insertion breakpoint. For each of the six carriers we separated the Alu-bearing and non-Alu (reference) haplotypes and computed pairwise distances across the window as the normalised Hamming distance (1 - proportion of shared alleles), visualising the relationships by hierarchical clustering (average linkage).

All analyses were performed using custom Python or R scripts and are included in our GitHub repository (data availability).

## Results

### Landscape of Alu and LINE1 elements in Indian populations

By benchmarking our short-read configuration against our overlapping long-read gold standard, we evaluated the performance of MELT-SPLIT across all retrotransposition-competent element families: Alu, LINE1, SVA, and HERV-K. Both Alu and LINE1 insertion calls achieved high precision (>0.80; Table S2), with corresponding mean F1 scores of 0.67 and 0.56, respectively (recall of 0.54 and 0.43 respectively). Conversely, SVA and HERV-K insertion calls were few in number, with lower precision (0.74 and 0.78, respectively) and were fully excluded from subsequent analyses.

We identified 7,075 unique LINE1 and 26,977 unique Alu insertions. On average, each individual carries 107.51 non-reference LINE1 and 943.72 non-reference Alu elements, covering 241,875 and 251,660 bases in the genome respectively (Figure 1A). The allele frequency distributions for MEIs are heavily right-skewed, with both Alus and LINE1s having a median alternate allele frequency of 0.001 (Figure S2). Alu insertions are primarily full-length, whereas LINE1 insertions show 5’-truncation (Figure S3). MEI length did not correlate with allele frequency for common LINE1s (MAF > 0.01; p=0.06) or common Alus (MAF>0.01; p=0.2) (Figure S4).

**Figure 1.**
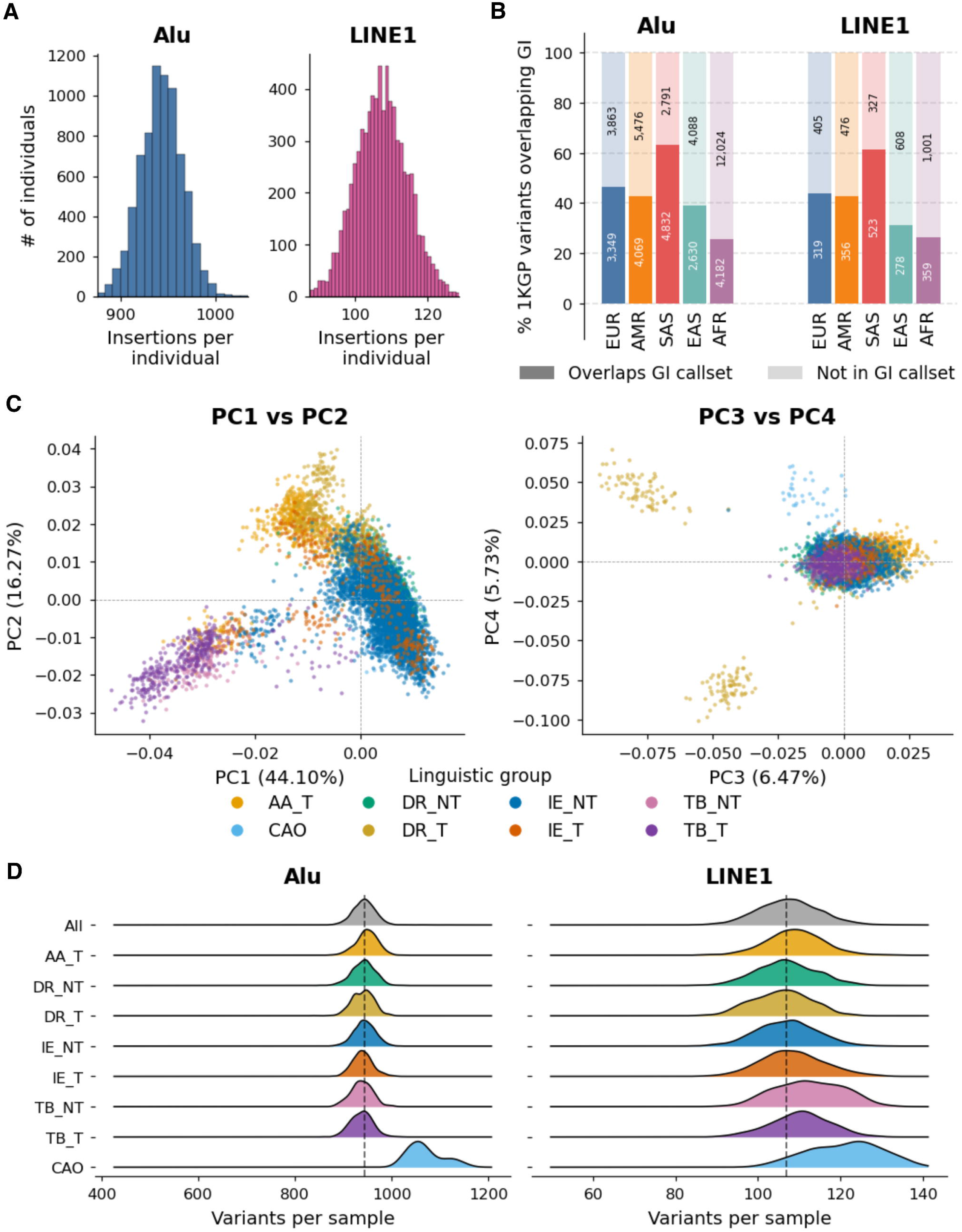
Landscape of MEIs in GI. A. Distribution of non-reference LINE1 and Alu insertions in GI. B. Percent of MEIs in the five 1000Genomes super populations in the GI dataset C. Principal component analysis (PCA) plots derived strictly from common MEI genotypes, plotting PC1 vs PC2 (left) and PC3 vs PC4 (right), color-coded by macro-population linguistic groups. D. Number of non-reference MEIs carried by each population group in GI;dotted line represents overall median

18,982 discovered MEIs are previously-unreported in other datasets; of these, 5,758 represent stable, non-singleton segregating variants (Table 1, Table S4). Of all variants in GI, 797 LINE1s (11.26%) and 7,447 Alus (27.59%) were found in the 30X 1000Genomes MEI dataset generated by Byrska-Bishop *et al.*, 2022, with maximum overlap expectedly with the South Asian subset of the dataset (Alus: 63.4%; LINE1s: 61.6%); this is despite South Asians making up a small fraction of the 1000Genomes dataset (Figure 1B).

Our dataset represents not one single Indian population, but 82 different ethnic groups. Based on the principal language families, we have previously classified the populations into eight population groups using a combination of linguistic family and tribal status. These are: tribal and non-tribal groups from the Indo-European (IE), Dravidian (DR), Tibeto-Burman (TB) families, and tribal groups from the Austroasiatic (AA) family, and an additional continentally admixed outgroup (CAO) representing a population known to be of admixed African descent^11^ (Table S1). A principal component analysis (PCA) of identified common (MAF > 0.01) MEIs recapitulated broad population structure in GI, particularly the NorthEast-west gradient (Figure 1C, Figure S5). PCs 3 and 4 also separated out the tribal populations from the Nilgiri hills, a population that also separates based on SNPs. On average, individuals in the CAO population carry more non-reference MEIs than other GI populations and groups (Figure 1D), which is also consistent with their distinct ancestry; this result also recapitulated the same patterns from SNP data.

### Genomic distribution of TEs

TE variants can disrupt genomic function if they are present in functionally relevant regions of the genome. Both Alu and LINE1 insertions were primarily found in intronic and intergenic regions, showing significant depletion in exons (Figure 2A, Table S5, Table S6); the magnitude of this depletion was stronger for common MEIs, suggesting that selection acts against insertions in functional, exonic regions.

**Figure 2.**
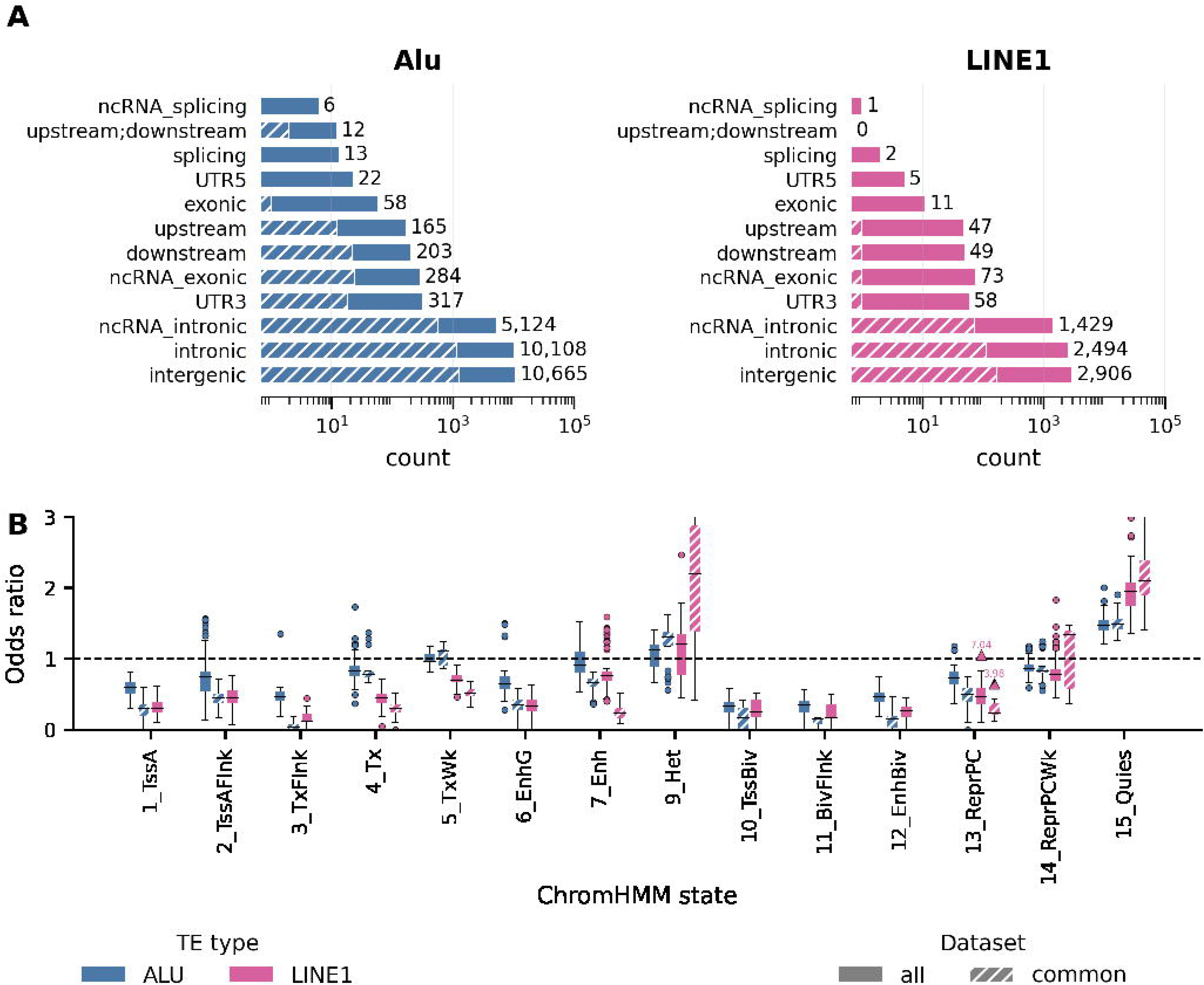
Pattern of MEIs across the genome. A. Genomic distribution of MEIs across ANNOVAR categories; hatched regions represent proportion of common MEIs in each category; numbers at the end represent total count of all MEIs in each category B. Enrichment of all and common MEIs across 15 Roadmap states, bars representing every tissue having an OR with pval≤ 0.05; dots represent outliers beyond 1.5×IQR, triangles represent extreme outliers beyond 1.5×IQR that do not fit within the axes represented in the figure.

Chromatin-state annotations across the 127 Roadmap epigenomes recapitulated this signal (Figure 2A; Table S7). Insertions were enriched in quiescent chromatin (15_Quies) in all 127 epigenomes (Alu OR ≈ 2.0, LINE1 OR ≈ 3.0) and depleted across active states, with LINE1 depleted more strongly than Alu in every tissue (median OR 0.63 vs 0.94; Wilcoxon V = 0, p < 2.2 × 10⁻¹⁶). Consistent with previous studies^9^, depletion in active chromatin was higher among common insertions while the quiescent enrichment was not, indicating that insertions accumulate in mutation-tolerant inactive chromatin, and are more likely to be removed when they land in active regulatory or transcribed regions. Among elements overlapping a regulatory state in ≥ 100 tissues (336 LINE1, 2,604 Alu) (Table S5), 38 non-singleton LINE1 and 358 non-singleton Alu insertions (of them 10 LINE1s and 137 Alus absent from other MEI databases) are private to a single population in GI and are thus candidate population-specific regulatory variants.

Both Alu and LINE1 insertions were depleted in early-replicating domains (ERD; Alu OR = 0.90, p = 4.77 × 10⁻¹⁷; LINE1 OR = 0.51, p = 2.00 × 10⁻^132^) and enriched in late-replicating domains (LRD; Alu OR = 1.50, p = 2.23 × 10⁻²⁴¹; LINE1 OR = 2.20, p = 2 × 10⁻^238^), with LINE1 showing the stronger effect in both directions (Table S8 for common MEIs).

Kojima et al. (2023)^28^ reported that replication timing bias is ancestry-dependent, with EAS (Japanese)-specific Alu insertions relatively enriched in early replicating domains (ERD) and non-EAS-specific insertions in late replicating domains (LRD). Given the genetic affinity of Tibeto-Burman (TB) populations to East Asian groups^11,29,30^, we tested this using group-specific Alu insertions in GI. Alus were enriched in LRDs in all four non-TB groups (CAO OR = 1.75; IE OR = 1.45; AA OR = 1.35; DR OR = 1.40; all p < 0.01), and depleted in ERDs in the two largest-effect groups (CAO OR = 0.83, p = 2.86 × 10⁻^3^; IE OR = 0.91, p = 0.017) (Table S9). Only TB-specific insertions showed no depletion in ERD (OR = 1.02, p = 0.83) and the weakest, non-significant LRD enrichment (OR = 1.22, 95% CI 0.99–1.48, p = 0.052; n = 400). This is unlikely to be a power artifact: the smaller AA set (n = 290) had significant LRD enrichment. The attenuated pattern in the EAS-adjacent TB group is directionally consistent with Kojima et al., though not significant. To test whether this attenuation reflects ancestry rather than sampling, we compared TB against the pooled non-TB groups by Cochran’s Q. Neither domain showed significant heterogeneity (ERD Q(1) = 1.14, p = 0.28; LRD Q(1) = 3.54, p = 0.06). TB-specific insertions are therefore directionally consistent with the pattern Kojima et al. report for EAS-specific insertions, but are not statistically distinguishable from the other GI groups at the current sample size (n = 400 TB-specific insertions).

### European-defined TE regulatory variants are often absent or differently distributed in Indian populations

Beyond their genome-wide distribution, TEs carry regulatory variation that is structured by ancestry, implying their function cannot be captured by European reference resources. We saw this along three lines: regulatory variants that Indian populations lack entirely, variants with sharply different frequencies between different Indian ethnicities, and variants that tag known trait associations.

A substantial fraction of European-defined regulatory MEIs are absent in GI. Of 1,670 Alu and LINE1 insertion eQTLs catalogued in GTEx (a primarily European-ancestry resource), 382 (23%) had no corresponding MEI call in GI; out of an independent set of 172 TE-eQTLs from 1000 Genomes LCLs, 67 (39%) were absent in GI. Low recall could account for some of these, but many likely mark loci where the European-defined regulatory variant is not segregating in Indian populations. This interpretation is supported by the fact that 222 of the 346 GTEx-absent Alu-eQTLs (64%) are also absent in IndiGen, arguing against under-calling as the sole explanation.

Even among those MEI-eQTLs that are present in GI, allele frequencies often vary widely across populations (Figure 3A, Figure S6), such that a regulatory variant common in one population can be near-absent in another. For example, a 1,486-bp LINE1 insertion in GI (chr8:130028989) has been previously shown to be an eQTL for the gene *CYRIB* in the thyroid, and lies in a known enhancer of *CYRIB* (Figure 3B). This insertion has an AF of 0.24 in GI and 0.18 in 1000G SAS; however it reaches an AF of 0.64 in TB_NER_1_01, and has a median AF of 0.51 across all TB populations, indicating an increase in the ‘downregulating’ allele in TB populations relative to other parts of the country. CYRIB binds active GTP-bound RAC1 through its switch-I loop, and inhibits RAC1-driven cytoskeletal remodeling^31–33^. The hyperactive RAC1 splice variant RAC1b, which shares the switch-I interface contacted by CYRIB^34^, is overexpressed in ∼33% of follicular thyroid carcinomas ^35^ and in BRAF V600E-positive papillary thyroid carcinomas^36^. Whether CYRIB also directly binds and inhibits RAC1b has not been tested, but the shared binding interface suggests this is plausible. A LINE1 insertion that constitutively reduces *CYRIB* expression would be predicted to disinhibit RAC1 signaling in thyroid, potentially promoting proliferation and invasive capacity, thus having clinical implications. 646 such eQTLs are strongly population-biased (allele frequency range > 0.3) within GI (Table S14), extending the set of genes carrying signals of differential regulation across Indian populations.

**Figure 3.**
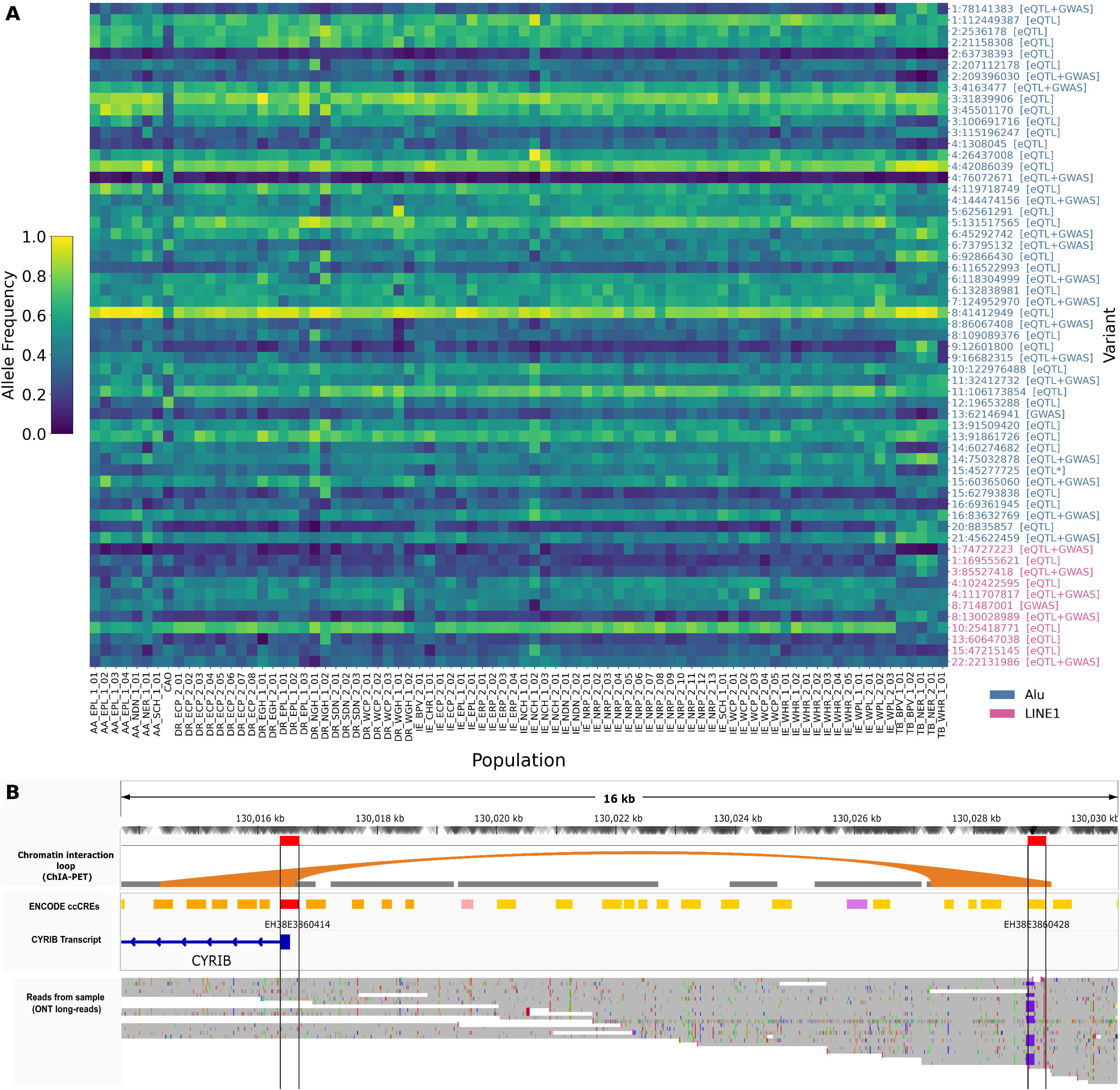
Functional regulatory MEIs in GI. A. Variation of allele frequencies (AF) of a subset of selected functional MEIs with an AF range>=0.4 across 82 population groups in GI; eQTL indicates an MEI-eQTL, that may also tag an eQTL SNP; eQTL* indicates an MEI that tags an eQTL SNP but is not documented to be an MEI-eQTL; GWAS indicates an MEI that tags a GWAS SNP B. LINE1 insertion which is an eQTL of the *CYRIB* gene lies in a distal enhancer of *CYRIB*; enhancer region and gene promoter interact through CHiA-PET interactions; insertions are shown in long-reads as purple markers

MEIs in high LD with trait-associated variants are themselves candidate functional alleles, either directly, or by altering the local regulatory landscape^37^. 12 common LINE1 and 119 common Alu insertions were in LD (r^2^>0.7) with at least one GWAS SNP (Table S10). Similarly, 18 common LINE1 and 140 common Alu insertions were in LD with a known eQTL SNP (Table S11). These regulatory-SNP tagging variants also showed a strong population bias within GI (Figure 3A).

A 280-bp intronic Alu insertion (chr16:69361945) both tags an eQTL SNP, and is also a reported MEI-eQTL for the gene *TERF2* (OMIM:602027)^26^ (Figure S7), and is linked to downregulating the gene’s expression. *TERF2* is responsible for stabilizing T-loop formation in telomeres which serves as protection against inappropriate recognition of chromosome ends as double-strand breaks and subsequent end-to-end chromosomal fusion^38,39^. TERF2 also has a high haploinsufficiency score (0.87)^40^, and the severity of TERF2 dysfunction has been shown to dictate cell fate, with mild dysfunction promoting senescence and severe dysfunction promoting apoptosis^41^. This Alu insertion ranges from an allele frequency of 0.101 in population AA_NER_1_01 to 0.625 in population IE_NCH_1_03. Differing frequencies of this eQTL TE implies differing regulation of expression of dosage-sensitive *TERF2*, which could potentially lead to differing clinical phenotypes.

Across these three lines of evidence, GI contains several hundred candidate functional variants at established trait loci, and these are priorities for follow-up in South Asian cohorts. Together, absence, frequency divergence, and trait-tagging point to a layer of TE regulatory variation that is population-specific in India and largely invisible to the reference resources human genetics currently relies on.

### Exonic MEIs segregating in GI populations reveal distinct coding variants and founder effects

Sixty-nine MEIs fell within exons (11 LINE1, 58 Alu; Table S12), and 19 were non-singletons (Table 2, Table S13); these represent function-altering insertions that don’t merely arise in Indian populations but also segregate in them.

**Table 2.** All non-singleton exonic insertions identified in GI; number of populations from which the carriers originate, and checked for presence in eight MEI databases.

| TE Type | SV length | Insertion Coordinate | Gene | Carriers <sup>b</sup> | # Populations | Unreported | LOEUF score for gene <sup>43</sup> |
| --- | --- | --- | --- | --- | --- | --- | --- |
| Alu | 281 | chr1:120810467 | *NBPF26* | 3 | 3 | Yes | 1.57 |
|  | 281 | chr1:12881899 | *PRAMEF4* | 610 | 79 | No | 0.95 |
|  | 273 | chr2:11174985 | *SLC66A3* | 4 | 4 | No | 1.6 |
|  | 276 | chr3:46265884 | *CCR3* | 2 | 1 | Yes | 1.93 |
|  | 279 | chr4:127811625 | *HSPA4L* | 2 | 1 | Yes | 0.55 |
|  | 245 | chr5:79737711 | *CMYA5* | 25 (1)<br>+1 related carrier | 14 | No | 0.81 |
|  | 87 | chr8:100573918 | *SNX31* | 14<br>+1 related carrier | 8 | No | 1.1 |
|  | 280 | chr9:7798574 | *DMAC1* | 2 | 1 | Yes | 1.84 |
|  | 281 | chr10:94706873 | *CYP2C18* | 5 | 1 | No | 1.16 |
|  | 281 | chr11:7695684 | *OVCH2* | 65 (1) | 28 | No | 1.51 |
|  | 279 | chr11:89714197 | *TRIM77* | 30 | 10 | No | 1.14 |
|  | 281 | chr11:113780379 | *CLDN25* | 5 | 5 | No | 1.89 |
|  | 279 | chr18:6850866 | *ARHGAP28* | 2 | 1 | No | 0.8 |
|  | 248 | chr18:24159972 | *CABYR* | 5 +1 related carrier | 1 | Yes | 1.18 |
|  | 279 | chr19:37699233 | *ZNF607* | 2 | 1 | Yes | 0.97 |
|  | 279 | chr19:38808903 | *LGALS4* | 2 | 1 | Yes | 1.18 |
|  | 280 | chr21:32269045 | *MIS18A* | 3 | 1 | Yes | 1.37 |
| LINE1 | 2066 | chr1:59915947 | *CYP2J2* | 12 | 5 | No | 1.06 |
|  | >4246 | chr4:99279553 | *ADH1A* | 2 | 2 | Yes | 1.1 |

<sup>b</sup> number of homozygous carriers

The clearest case is a LINE1 insertion in exon 2 of gene *CYP2J2* (OMIM:601258)^26^ at position chr1:59915947 (Figure 4A), a cytochrome P450 that epoxidizes arachidonic acid in cardiac tissue and whose coding variants are tied to cardiovascular disease and Alzheimer’s Disease. Twelve individuals from five populations (Figure 4B) are heterozygous carriers for this insertion. The variant has also been previously found in a single South Asian individual in 1000Genomes and one non-Finnish European individual in gnomAD, but not characterized.

**Figure 4.**
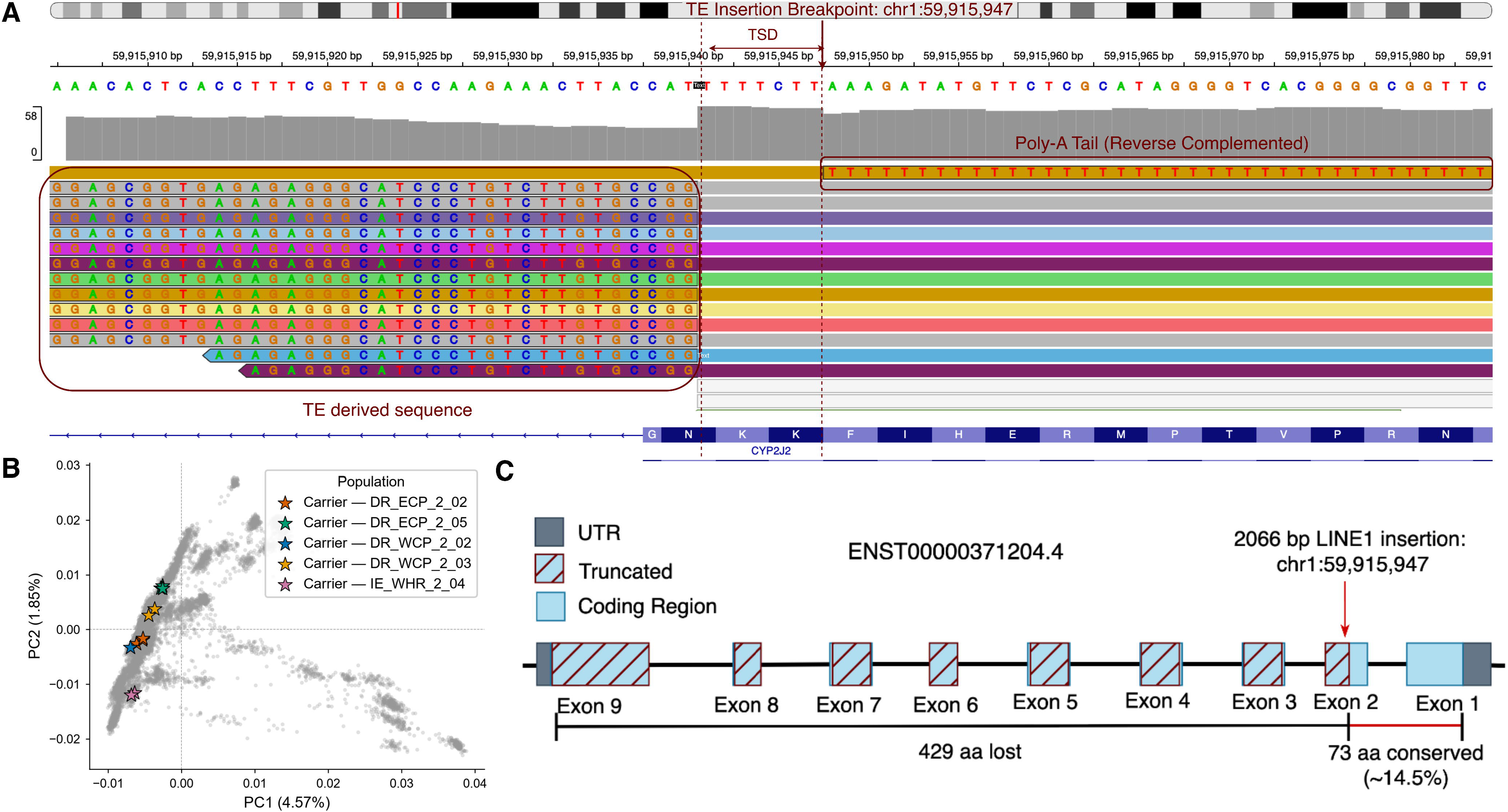
Exonic insertion in gene *CYP2J2.* A. IGV plot showing the reads supporting the insertion from long-read sequencing data in one carrier. B. Carriers of the heterozygous insertion in gene *CYP2J2* spread in GI C. Location of the insertion in the gene region.

Exon 2 is retained in most *CYP2J2* isoforms, so the insertion plausibly yields a truncated protein (Figure 4C). Its pHaplo score is moderate (0.69)^40^, so heterozygote carriers can potentially remain unaffected. Indeed, carriers of this mutation do not show significant differences in lipid levels or increased likelihood of disease compared to non-carriers from the same community.

Its enrichment in specific communities (AF > 1% in two Dravidian non-tribal communities) increases the likelihood of homozygosity for the insertion, leading to the possibility of increased knockout and thus disease risk, and making this a clear target for functional follow-up. The variant has been confirmed by long-read sequencing in seven of these carrier samples (Figure S8).

A second, previously unreported variant, a 248-bp Alu insertion (chr18:24159972) in *CABYR* (Figure S9A), reached an allele frequency of 0.035 within unrelated individuals of a single endogamous community (DR_NGH_1_01, Figure S9B), absent elsewhere. Moreover, one parent-child pair carries the insertion, confirming its Mendelian inheritance. The haplotypes carrying the insertion cluster together compared to other haplotypes in the carrier samples, and are longer (Figure S10B), indicating a founder effect.

*CABYR* encodes a testis-specific protein required for sperm capacitation, with prior links to reduced motility. The insertion overlaps exon 5 (the terminal coding exon) of the MANE select transcript (which is also the highest expressed isoform in the testis). In this transcript, the insertion falls in codon 348 of 379, within the terminal coding exon; the inserted Alu sequence leads to a premature termination codon 285 nt upstream of the final exon–exon junction and satisfying the threshold for nonsense-mediated mRNA decay. We thus predict loss of function from this allele for transcripts of this architecture.

However, *CABYR* encodes multiple functionally distinct isoforms, and the consequence of this insertion is transcript-dependent. In the other isoform (longer, 493 amino acids), the insertion lies in the 3’ UTR. The overall impact of the insertion on CABYR function, therefore, needs experimental validation. Testing the consequences of this mutation on fertility would be a future direction. A coding-exon variant which is common in one community, and invisible across the rest of the cohort, is the kind of private allele a study like GI is built to find.

## Discussion

This study represents a large characterization of mobile element variation in diverse Indian populations, spanning 82 populations and including geographically isolated and previously unreported communities. We demonstrate the presence of large untapped genetic variation at transposable elements, and report 18,982 previously-unreported variants (5,758 non-singletons) across the genome. Beyond cataloging this variation, we also report strong variation between populations; a substantial fraction of TE regulatory variation is either absent from, or carried at markedly different frequencies, across Indian communities relative to European-biased reference resources. Insertions bear signatures of purifying selection, with depletion in exons and active regions of the genome, and enrichment in quiescent regions.

A meaningful fraction of regulatory variants identified in European populations do not segregate in Indian populations, while others have different frequencies in different Indian populations. These point to a layer of population-specific regulatory variation that reference resources built on European-ancestry cohorts do not capture. Because this variation is population-structured, MEIs are a plausible and largely unexamined contributor to the reduced transferability of genetic associations across ancestries. Testing that contribution directly, however, will require functional genomic resources (including expression, methylation, and chromatin data), generated in South Asian populations rather than transferred from European-ancestry references. Building those resources is the necessary complement to catalogs such as this one.

The presence of exonic MEIs strongly supports the role of MEIs in population-specific gene function, and highlights its potential clinical relevance. Nineteen of these non-singleton insertions lie within exons, of which nine haven’t been reported previously. Of these, the LINE1 insertion in *CYP2J2* illustrates how a mobile element can potentially disrupt a clinically relevant gene. The insertion falls in exon 2, retained in most *CYP2J2* isoforms, and is predicted to yield a truncated product. *CYP2J2* encodes a cytochrome P450 that generates epoxyeicosatrienoic acids in cardiac tissue, and its coding variation has been linked to cardiovascular disease. This insertion exceeding 1% frequency in two Dravidian communities, while appearing only in a single individual earlier, underscores the central point of this study: functionally consequential coding variation can be common within Indian communities while remaining nearly invisible to existing resources. This population-enriched candidate warrants targeted functional and clinical follow-up.

While historical founder effects are well-documented drivers of rare recessive disease prevalence in highly endogamous Indian communities, our data show that transposable elements can follow the same evolutionary trajectory. The localized, high-frequency *Alu* insertion in *CABYR* within a single tribal community serves as a clear paradigm for how population-specific endogamy can rapidly elevate potentially deleterious structural variants to high frequencies.

Our study has important limitations, primarily stemming from the short-read datasets used. The modest recall for TEs makes this callset an undercount of the true variants segregating in the population. Thus, absence in our callset reflects detection as well as biology. This challenge may be particularly high for genotype-level inference at individual loci, and has led to the exclusion of deletions from our callset. Moreover, comparisons drawn against European-ancestry GWAS and eQTL resources are necessarily indirect, given the variation in breakpoints called by different callers in different datasets. Long-read sequencing at population scale will resolve much of these limitations, yielding more complete and more accurately genotyped MEI callsets. This work maps where in India’s populations that fuller picture is most likely to be informative.

## Supporting information

Consortium Author List

## Supplemental Information

Supplemental information consists of 10 figures, 14 tables and 1 supplemental text.

Document S1: Figures S1-S10, Tables S8 and S9, and Text S1

Data S1: Supplemental Tables S1-S7, S10-S14

## Acknowledgements

We thank all participants who consented to providing their samples for this project. We acknowledge the funding by the Department of Biotechnology (DBT), Ministry of Science and Technology, Government of India. We thank the CBR BioBank for help accessing samples for validation, and the CCMB sequencing core for generating long-read sequencing data. We acknowledge Anand Kumar and Jothibasu from CBR’s HPC team for compute resources, and Bratati Kahali for data sharing and help with the manuscript.

## Data and Code Availability

Raw data are available from the Indian Biological Data Centre (IBDC) upon request under IBDC’s FeED protocol. All identified MEIs, including population-specific allele frequencies, are included in the Supplementary Table S14.

All code used in this analysis is available at https://github.com/gsshreya/TE_analysis/, as well as https://doi.org/10.5281/zenodo.21615044

## Ethics statement

The GenomeIndia consortium adhered to the principles of the Helsinki Declaration for research protocols, consent forms, sample collection, and ethical practices. These guidelines were uniformly implemented across all sampling centres and approved by their respective institutional human ethics committees (IECs). A detailed ethics statement is mentioned in Subramanian et al 2026^11^.

## Web Resources

Public datasets used in this study are available at:

1KGP Phase 3: https://ftp.1000genomes.ebi.ac.uk/vol1/ftp/phase3/integrated_sv_map/supporting/GRCh38_positions/ALL.wgs.mergedSV.v8.20130502.svs.genotypes.GRCh38.vcf.gz

1KGP Phase 4: https://ftp.1000genomes.ebi.ac.uk/vol1/ftp/data_collections/1000G_2504_high_coverage/working/20210124.SV_Illumina_Integration/1KGP_3202.gatksv_svtools_novelins.freeze_V3.wAF.vcf.gz

1KGP ONT-Vienna longreads: https://ftp.1000genomes.ebi.ac.uk/vol1/ftp/data_collections/1KG_ONT_VIENNA/release/v1.1/svan-annotation/final-vcf.unphased.SVAN_1.3.vcf.gz

gnomAD SV v4: https://storage.googleapis.com/gcp-public-data--gnomad/release/4.1/genome_sv/gnomad.v4.1.sv.sites.vcf.gz

SGDP: https://academic.oup.com/gbe/article/12/6/779/5828221#supplementary-data

nstd211: https://www.ncbi.nlm.nih.gov/dbvar/studies/nstd211/download/?type=i

HMEID: http://bigdata.ibp.ac.cn/HMEID/download/file/MEI.GRCh38.HMEIDv1.1.vcf.gz/

Indigen: https://clingen.igib.res.in/indigen/download/Indigen_Alu_final_geno10_all_22K.vcf

Epigenomics Roadmap: https://egg2.wustl.edu/roadmap/data/byFileType/chromhmmSegmentations/ChmmModels/coreMarks/jointModel/final/download/

Replication-timing Annotations: https://ftp.ncbi.nlm.nih.gov/geo/series/GSE53nnn/GSE53984/suppl/GSE53984%5FGSM923453%5FBg02es%5FRep1%5Fsegments%2Ebed%2Egz

NHGRI-EBI GWAS catalogue (all_associations v1.0): https://www.ebi.ac.uk/gwas/docs/file-downloads

Adult GTEx Single-Tissue cis-QTL data (V11): https://storage.googleapis.com/adult-gtex/bulk-qtl/v11/single-tissue-cis-qtl/GTEx_Analysis_v11_eQTL.tar

GTEx MEI eQTLs: https://pmc.ncbi.nlm.nih.gov/articles/instance/7385971/bin/13059_2020_2101_MOESM7_ESM.txt

1000G LCL MEI eQTLs: https://static-content.springer.com/esm/art%3A10.1186%2Fs12859-019-3113-x/MediaObjects/12859_2019_3113_MOESM3_ESM.xls

gnomAD web browser: https://gnomad.broadinstitute.org/

IGV (Integrative Genomics Viewer): https://igv.org/

Segmental Duplication Data: https://hgdownload.soe.ucsc.edu/goldenPath/hg38/database/genomicSuperDups.txt.gz

ClinVar: https://www.ncbi.nlm.nih.gov/clinvar/

Tools used in this study are available at:

MELT v2.2.2: https://melt.igs.umaryland.edu/downloads.php

BCFTools v1.21: https://github.com/samtools/bcftools/releases/#release-1.21

Dorado v0.5.2: https://github.com/nanoporetech/dorado

xTea_long_release_v0.1.0: https://github.com/parklab/xTEA

Sniffles v2.3.3: https://github.com/fritzsedlazeck/sniffles

ANNOVAR: https://annovar.openbioinformatics.org/en/latest/user-guide/download/

PLINKv1.9:https://www.cog-genomics.org/plink/

PLINKv2: https://www.cog-genomics.org/plink/2.0/

