## Supplemental Figures and Text for "Population-Scale Landscape of Mobile Element Insertions Across 82 Diverse Indian Populations"

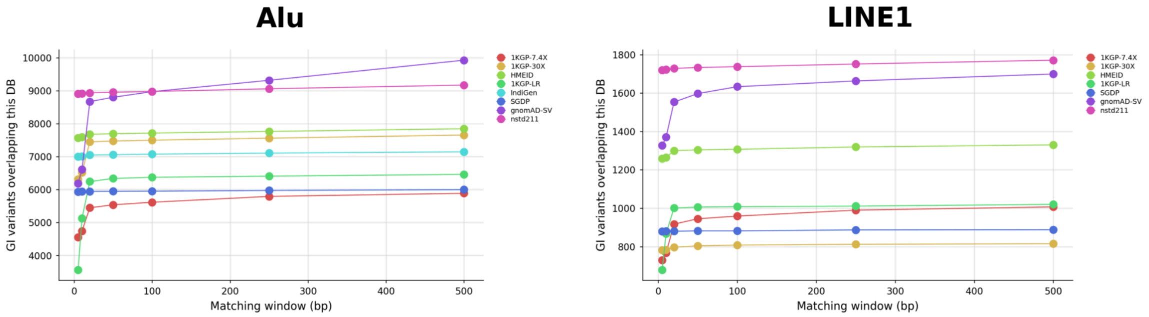


**Figure S1**. Overlap in Alus and LINE1 insertions with other databases as a function of window size used for defining overlap. Window is defined as x bp upstream and downstream of the insertion.

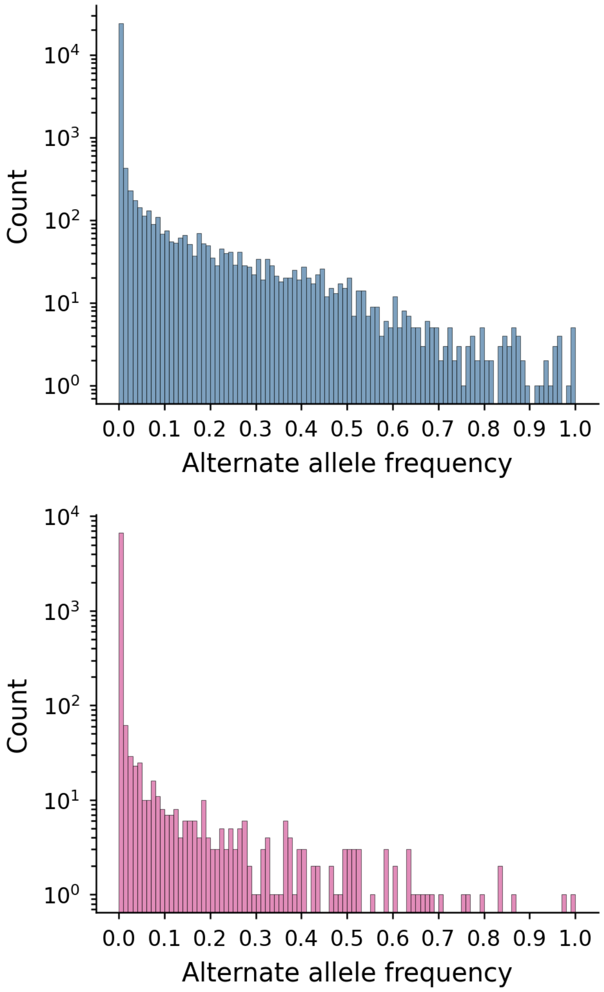


**Figure S2** Allele frequency spectrum of Alu (top) and LINE1 (below) MEIs in GI

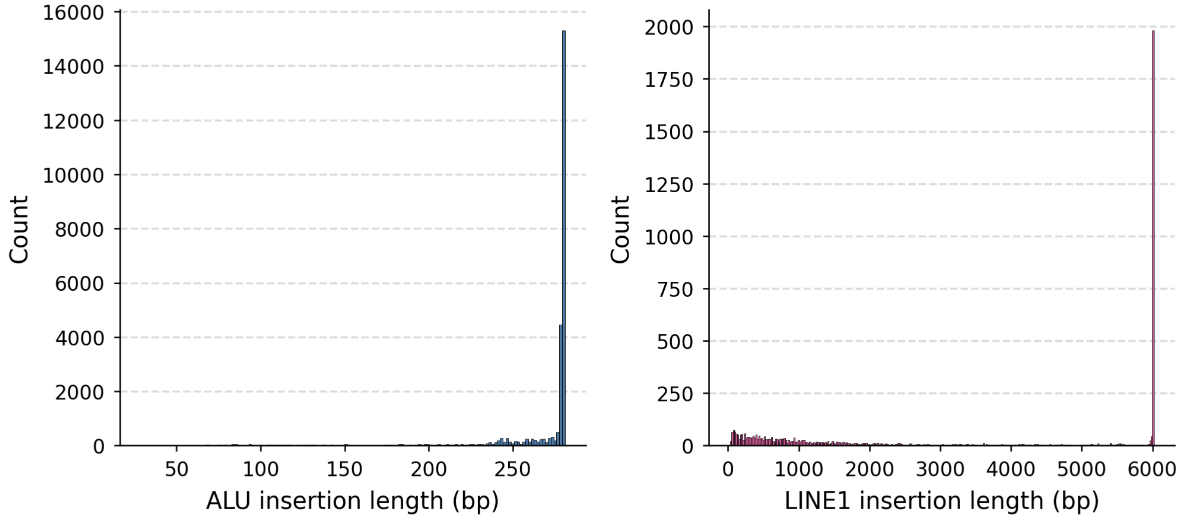


**Figure S3** Length distribution of Alu and LINE1 MEIs in GI

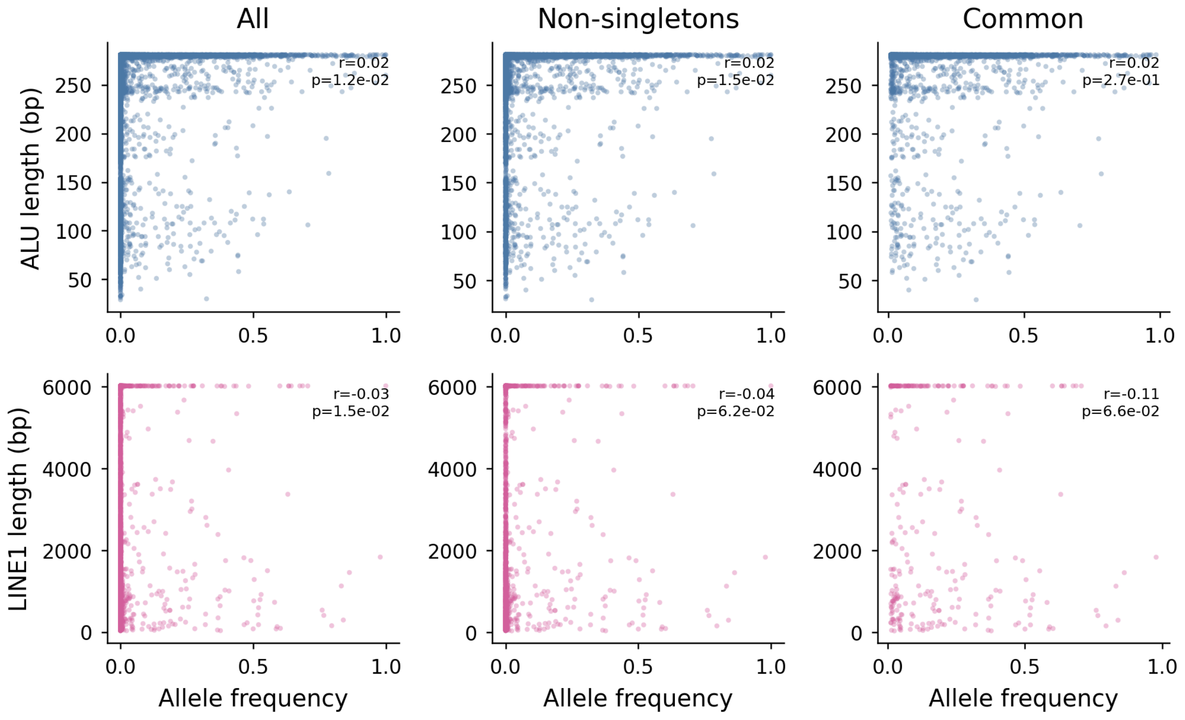


**Figure S4**. Correlation between length and allele frequency for Alus (top) and LINE1s (bottom)

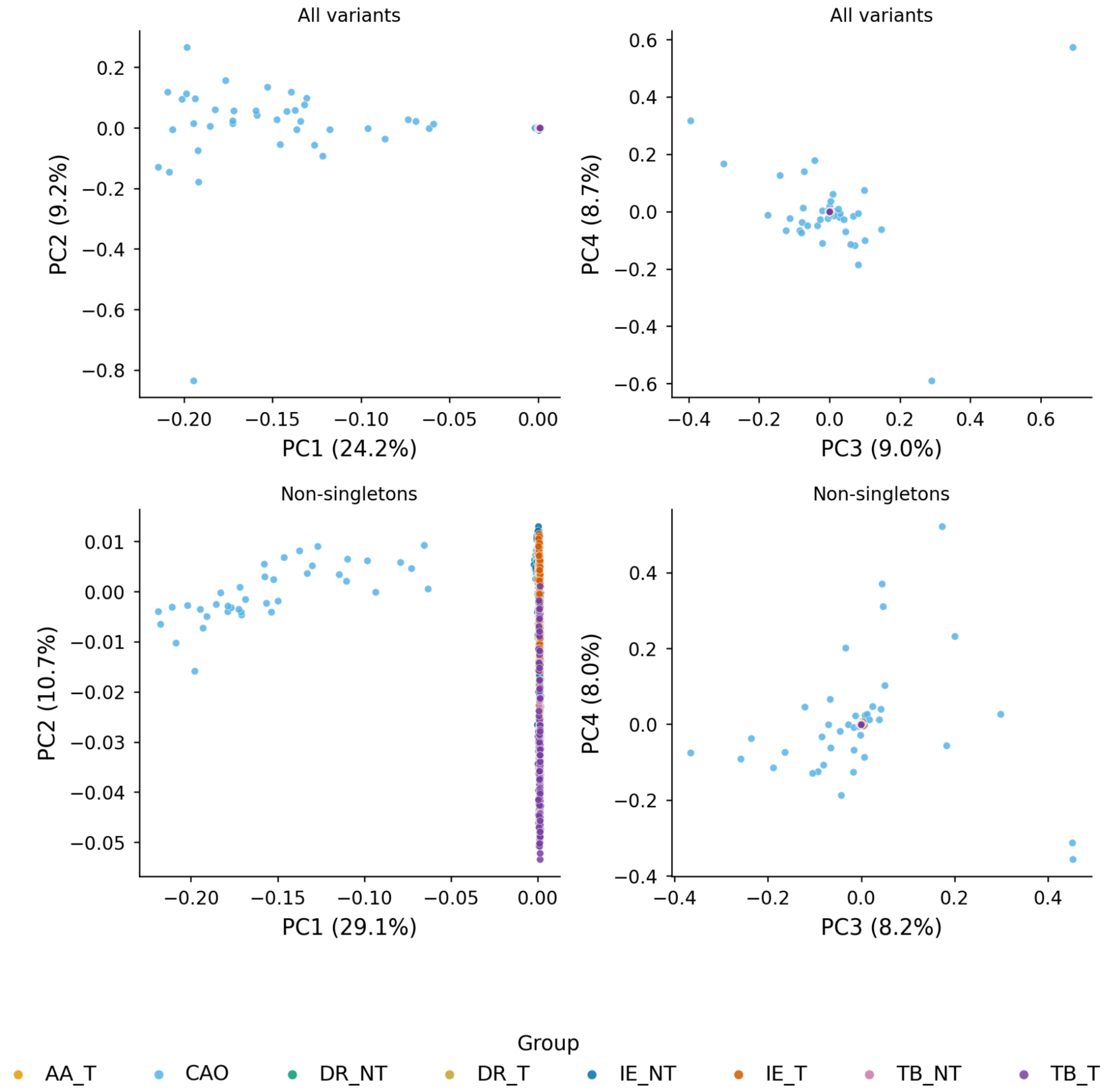


**Figure S5.** PCA using all MEIs (top) and non-singleton MEIs (>=2 carriers, bottom) in GI

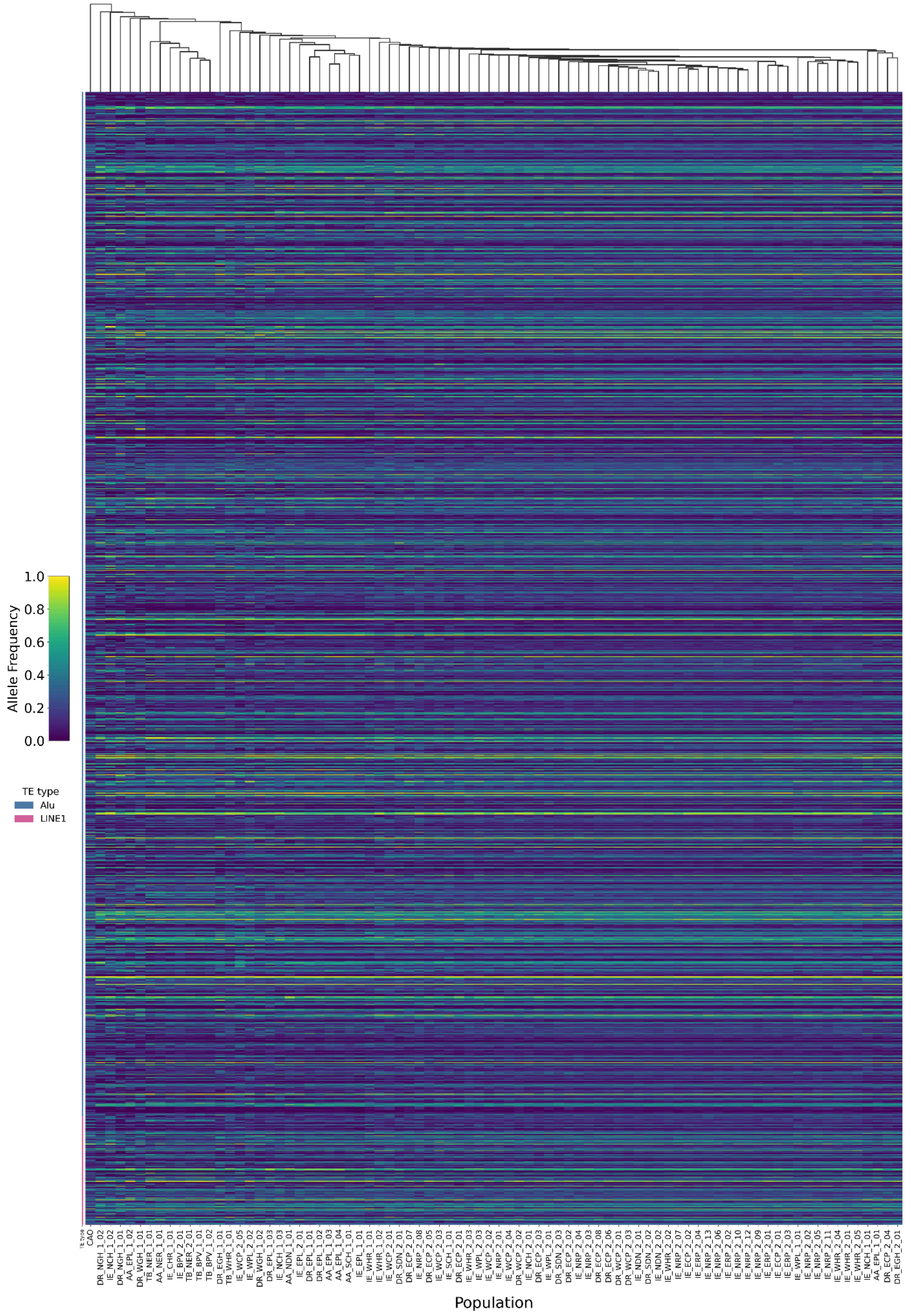


**Figure S6**. Heatmap of allele frequencies of all functional variants (MEI eQTLS, MEIs tagging eQTLS, MEIs tagging GWAS SNPs) in GI, across 82 populations (clustered)

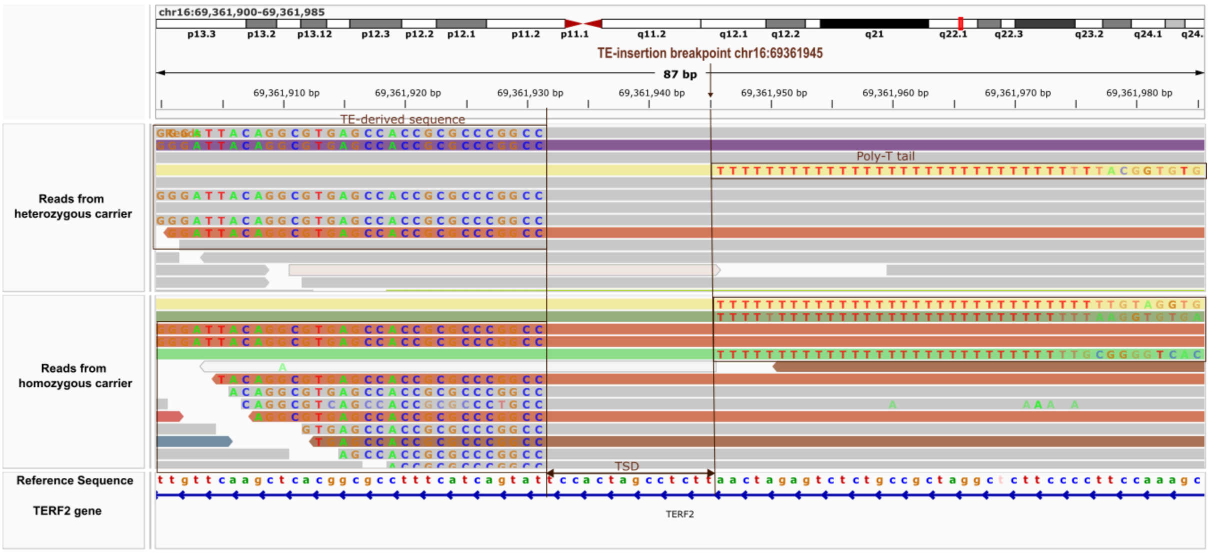


**Figure S7**. Alu-insertion at chr16:69361945 in the intron of *TERF2* gene, that acts as an eQTL of TERF2

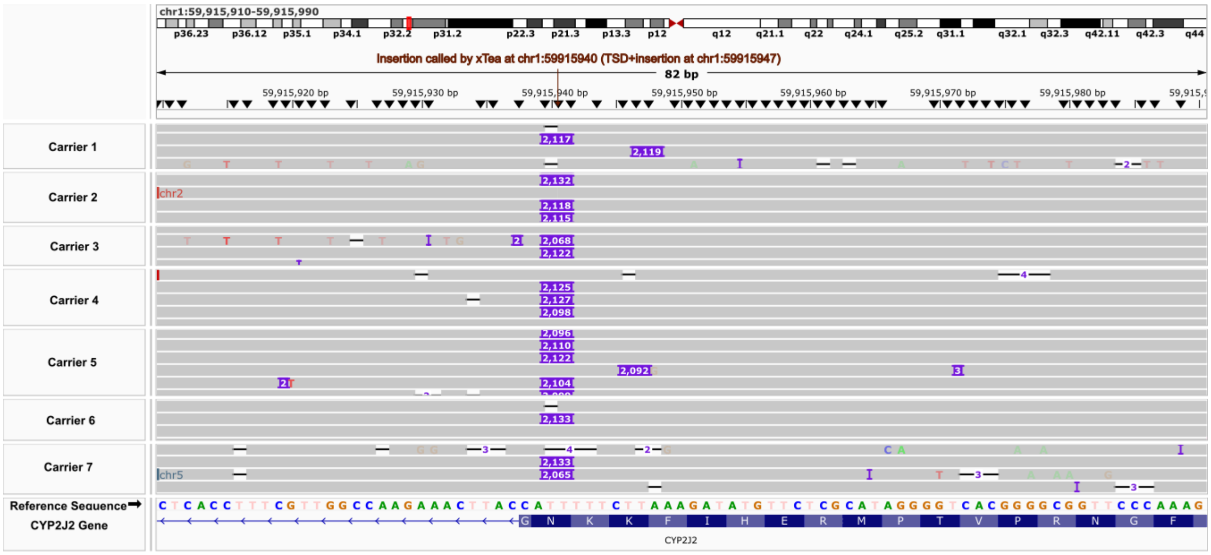


**Figure S8**: IGV plot showing reads supporting insertion in *CYP2J2* from long read sequencing data in seven carriers

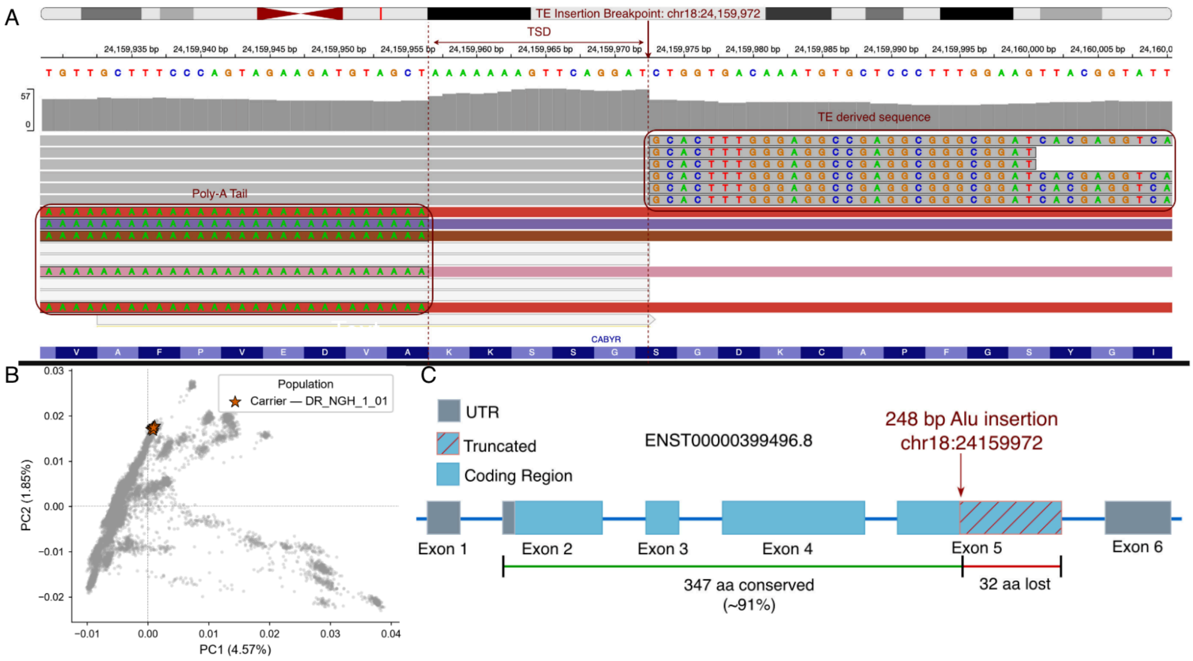


**Figure S9.** Exonic insertion in *CABYR*. A. IGV plot showing the reads supporting the insertion from short read sequencing data in one carrier. B. Carriers of the heterozygous insertion in gene *CABYR* spread in GI C. Location of the insertion in the gene region.

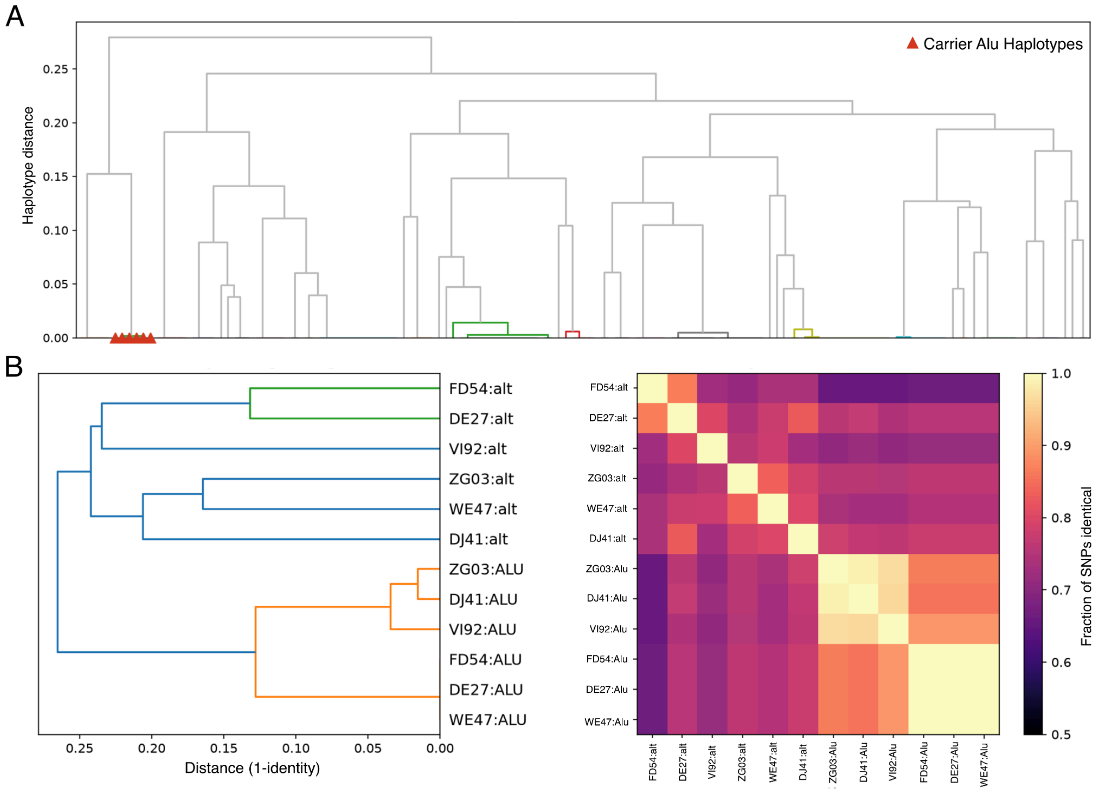


**Figure S10.** Haplotype analysis of the Alu insertion variant in the CABYR gene A**.** Population-matched, core block. The six Alu haplotypes (red triangles) form one tight cluster at ~zero distance, distinct from the rest of the population. B.Carrier Alu-bearing vs. non-Alu (reference) haplotype clustering across the ±500 kb window.

### Supplemental Tables (remaining large supplemental tables are in an attached excel spreadsheet)

**Table S8.** ERD and LRD enrichment analysis of MEIs

| Dataset | TE Type | Total Number of Variants | ERD Odds Ratio | ERD 95% CI | ERD pvalue | LRD Odds Ratio | LRD 95% CI | LRD pvalue |
|---|---|---|---|---|---|---|---|---|
| All | Alu | 26,977 | 0.897 | 0.874–0.920 | $4.77 \times 10^{-17}$ | 1.501 | 1.465–1.537 | $2.23 \times 10^{-241}$ |
| All | LINE1 | 7,075 | 0.51 | 0.481–0.540 | $2.00 \times 10^{-132}$ | 2.199 | 2.097–2.307 | $1.99 \times 10^{-238}$ |
| Common | Alu | 3029 | 0.922 | 0.854–0.995 | $3.69 \times 10^{-2}$ | 1.402 | 1.305–1.507 | $2.57 \times 10^{-20}$ |
| Common | LINE1 | 362 | 0.389 | 0.290–0.514 | $1.98 \times 10^{-13}$ | 2.123 | 1.715–2.632 | $1.40 \times 10^{-12}$ |

**Table S9.** ERD and LRD enrichment analysis of population-specific Alu insertions

| Dataset | Total Variants | ERD Odds Ratio | ERD 95% CI | ERD pvalue | LRD Odds Ratio | LRD 95% CI | LRD pvalue |
|---|---:|---:|---|---|---:|---|---|
| AA | 290 | 0.935 | 0.724–1.201 | 0.6209 | 1.353 | 1.066–1.716 | 0.0114 |
| CAO | 1199 | 0.83 | 0.732–0.940 | $2.86 \times 10^{-3}$ | 1.749 | 1.559–1.963 | $5.00 \times 10^{-22}$ |
| DR | 1306 | 0.968 | 0.861–1.088 | 0.5998 | 1.404 | 1.258–1.568 | $1.20 \times 10^{-9}$ |
| IE | 2887 | 0.909 | 0.840–0.983 | 0.0167 | 1.451 | 1.348–1.562 | $2.75 \times 10^{-23}$ |
| TB | 400 | 1.022 | 0.826–1.260 | 0.8333 | 1.221 | 0.996–1.494 | 0.0515 |

### Supplemental Text

#### Text S1: Calling and benchmarking reference transposable element deletions

We identified reference TE deletions using the MELT-Deletion module with the combined AluY and LINE1 reference bed file provided by MELT. We filtered the raw deletion calls using BCFtoolsv1.21 to exclude all deletion variants with >25% missingness. The filtered deletion calls were then benchmarked against the long-read sequencing data to assess the accuracy of MELT calls.

Long-read sequencing samples (obtained as described in the methods of the main paper) were also used to benchmark deletions. We overlapped deletions reported by Sniffles with reference TE .bed files provided in the MELT .jar, and identified any deletions that had start and end positions within 50 bp of a reference TEs start and end positions as reference TE deletions. These reference TE deletions were used to benchmark MELT-Deletion calls, requiring ≥ 80% reciprocal overlap between MELT and Sniffles calls.

For deletions, though MELT-DELETION demonstrated high precision and F1 scores for Alus and LINE1s (precision of 0.96 and 0.90 respectively, and F1 of 0.89 and 0.8 respectively) (Table S2), it demonstrated a high error rate in separating heterozygous from homozygous calls . Because accurate genotype frequencies are foundational to downstream population structure and allele-sharing analyses, all subsequent functional and population genetic investigations were restricted strictly to TE insertions.
